# Milk Exosomes Cross the Blood-Brain Barrier in Murine Cerebral Cortex Endothelial Cells and Promote Dendritic Complexity in the Hippocampus and Brain Function in C57BL/6J Mice

**DOI:** 10.1101/2021.11.24.469889

**Authors:** Fang Zhou, Pearl Ebea, Ezra Mutai, Sonal Sukreet, Shya Navazesh, Haluk Dogan, Wenhao Li, Juan Cui, Peng Ji, Denise M. O. Ramirez, Janos Zempleni

**Affiliations:** Departmentsof Nutrition & Health Sciences, University of Nebraska-Lincoln, Lincoln, NE, USA; Department of Nutrition, University of California Davis, Davis, CA, USA; Department of Computer Science & Engineering, University of Nebraska-Lincoln, Lincoln, NE, USA; Department of Neurology, University of Texas Southwestern Medical Center, Dallas, TX, USA; Peter O’Donnell, Jr. Brain Institute, University of Texas Southwestern Medical Center, Dallas, TX, USA

**Keywords:** blood brain barrier, gene expression, milk exosomes, neuronal development, serial two-photon tomography

## Abstract

**Background:** Human milk contains large amounts of exosomes (MEs) and their regulatory microRNA cargos, whereas infant formulas contain only trace amounts of MEs and microRNAs. Breastfeeding has been implicated in optimal brain development but experimental evidence linking ME intake with brain development is limited.

**Objectives:** We assessed the transport of MEs across the blood-brain barrier (BBB) and ME accumulation in distinct regions of the brain in brain endothelial cells and suckling mice. We further assessed BME-dependent gene expression profiles and effects on the dendritic complexity of hippocampal granule cells and phenotypes of BME depletion in neonate, juvenile and adult mice.

**Methods:** The transfer of MEs across the BBB was assessed by using bovine MEs labeled with FM4-64 or loaded with IRDye-labeled miR-34a in murine brain endothelial bEnd.3 cell monolayers and dual chamber systems, and in wild-type newborn pups fostered to exosome and cargo tracking (ECT) dams that express MEs endogenously labeled with a CD63-eGFP fusion protein for subsequent analysis by serial two-photon tomography and staining with anti-eGFP antibodies. Effects of MEs on gene expression and dendritic architecture of granule cells was analyzed in hippocampi from juvenile mice fed exosome and RNA-depleted (ERD) and exosome and RNA-sufficient (ERS) diets by using RNA-sequencing analysis and Golgi-Cox staining followed by integrated neuronal tracing and morphological analysis of neuronal dendrites, respectively. Spatial learning and severity of kainic acid-induced seizures were assessed in mice fed ERD and ERS diets.

**Results:** bEnd.3 cells internalized MEs by using a saturable transport mechanism and secreted miR-34a across the basal membrane. MEs penetrated the entire brain in fostering experiments; major regions of accumulation included the hippocampus, cortex and cerebellum. Two hundred ninety-five genes were differentially expressed in hippocampi from male mice fed ERD and ERS diets; high-confidence gene networks included pathways implicated in axon guidance and calcium signaling. Only one gene was differentially expressed in females fed the experimental diets. Juvenile pups fed the ERD diet had reduced dendritic complexity of dentate granule cells in the hippocampus, scored nine-fold lower in the Barnes maze test of spatial learning and memory (*P* < 0.01), and the severity of seizures was 5-fold higher following kainic acid administration in adult mice fed the ERD diet compared to mice fed the ERS diet (*P* < 0.01).

**Conclusions:** MEs cross the BBB and contribute toward optimal neuronal development, spatial learning and memory, and resistance to kainic acid-induced seizures in mice.

## INTRODUCTION

Most cells synthesize and secrete nanoparticles called exosomes (∼100 nm) into the extracellular space [1]. Exosomes travel to adjacent and distant recipient cells and play an important role in cell-to-cell communication [1], including transfer across the blood-brain barrier (BBB^4^) [2; 3]. Communication is achieved through the transfer of regulatory exosome cargos from donor cells to recipient cells as well as binding of exosomes to receptors on the recipient cell surface [1; 4]. Among exosome cargos, microRNAs have gained particular attention because they regulate more than 60% of human genes and loss of microRNA biogenesis in Dicer knockout mice is embryonic lethal [5; 6]. MicroRNAs are short non-coding RNAs that bind to complementary sequences in the 3’-untranslated regions in mRNAs [7; 8]. If complementarity in the seed region (nucleotides 2-8 in microRNA) is perfect, mRNA is degraded [7; 8]; if complementarity is imperfect, mRNA translation is halted [9; 10].

We have pioneered a new line of discovery by demonstrating that exosomes and their microRNA cargos do not originate exclusively in endogenous synthesis but may also be absorbed from milk in human adults and neonate and adult mice and piglets [11; 12; 13]. Evidence is accumulating that endogenous synthesis of microRNAs cannot compensate for dietary depletion of exosomes and their microRNAs cargos. The concentrations of microRNAs were up to 60% lower in the plasma, liver, skeletal muscle, intestinal mucosa and placenta in mice fed an exosome and RNA-depleted (ERD) diet compared to controls fed an exosome and RNA-sufficient (ERS) diet [11; 14; 15; 16; 17]. The depletion of tissue microRNAs in mice fed ERD was associated with phenotypes such as altered purine metabolism, changes in bacterial communities in the gut, a moderate loss of muscle grip strength, increased severity of symptoms of inflammatory bowel disease and loss of fecundity and postnatal survival compared to ERS controls [14; 15; 16; 17; 18; 19]. Milk exosome (ME) supplementation studies reported an increase in villus height and crypt depth in the murine intestinal mucosa, reduced severity of inflammation in mouse models of necrotizing enterocolitis and improved bone health in mouse models of osteoporosis compared to non-supplemented controls [20; 21; 22].

These observations are of great importance in nutrition, particularly the nutrition of infants. The American Academy of Pediatrics recommends that human milk be the sole source of nutrition in the first six months of life [23]. Human milk contains large amounts of exosomes (2.2*x*10^11^/mL) loaded with more than 200 distinct microRNAs, whereas infant formulas are essentially free of MEs and microRNAs [24]. There may be implications of low ME intake for the optimal neurological development of infants. For example, white matter, sub-cortical gray matter volume and cortical thickness were greater in breastfed infants compared with formula-fed infants although cause-and-effect relationships between ME intake and brain development were not assessed [25]. Only 26% of parents in the U.S. fed their infants exclusively with human milk in the first six months of life in 2017 [23; 26]. The 2.8 million infants born annually in the U.S. that are partially or exclusively formula-fed do not realize the potential benefits conferred by MEs [26; 27].

Previously, we provided evidence that a large percentage of orally administered MEs and microRNAs cargos accumulate in the brain in suckling mice and piglets and adult mice [13]. These studies did not formally exclude the possibility that the MEs remained in the vasculature. In this paper we investigated the transport of MEs across the BBB and ME accumulation in distinct regions of the brain, ME-dependent gene expression profiles and functional effects such as neuronal development and brain phenotypes of ME depletion in cell culture models and mice. Studies of neuronal development focused on dendritic complexity because dendritic arborization and branching patterns are susceptible to modulation of environmental cues [28]. Exosomes are implicated in intercellular communication among neurons. For example, the injection of exosomes from neural cell cultures into the lateral ventricles of postnatal day 4 mice increased neural proliferation enhanced in dentate gyrus [29]. Motivated by this prior knowledge, we assessed the contribution of MEs toward optimal neuronal development and brain health in mice.

## METHODS

### Isolation and labeling of MEs

Bovine MEs (BMEs) were isolated from skim milk from a local grocery store by using sequential ultracentrifugation and authenticated by using Nanosight NS300 nanoparticle size analysis, scanning electron microscopy and transmission electron microscopy as previously described (**Supplemental Fig. S1**) [30]. The antibodies and their dilutions used in immunoblot analysis were the same as previously described [30]. Protocol details were deposited in the EV-Track database (ID EV210338). BMEs were suspended in sterile phosphate-buffered saline (PBS) and kept at −80°C until use. For transport studies in cell monolayers, BMEs were labeled with FM 4-64 (Molecular Probes, Inc.) or by labeling RNA cargos by using the ExoGlow-RNA^TM^ EV Labeling Kit (System Biosciences, Inc.) following the manufacturers’ recommendations. For transport studies in dual-chambers, BMEs were loaded with synthetic IRDye-labeled miR-34a as previously described [13].

### BME transport in cell cultures

Murine brain endothelial bEnd.3 cells [American Type Culture Collection (ATCC) CRL-2299, passages 21 - 30) and C8-D1A astrocytes (ATCC CRL-2541, passage unknown to ATCC) were purchased form ATCC. BV2 microglia (passage 15 - 25) were a gift from Dr. Sanjay Maggirwar (University of Rochester Medical Center, Rochester, NY, USA). Cells were cultured following ATCC recommendations. In monolayer studies, uptake of BMEs by bEnd.3 cells and BV2 microglia was assessed as previously described using times, concentrations and competitors shown in Results [Wolf, 2015 #11626]. Transport kinetics was modeled using the Michaelis-Menten equation and nonlinear regression; modeling was conducted using GraphPad Prism 6.0 (GraphPad Software). Confocal Z-stacks were collected at 60-fold magnification using 300 nm z-spacing on an A1R-Ti2 confocal system (Nikon) and used to determine whether bEnd.3 cells internalized BMEs or whether BMEs adsorbed to the cell surface. Dual chamber assays as a model of transport across the blood-brain barrier (BBB) were conducted as previously described with the following modifications [12]. bEnd.3 cells were seeded on the semiporous membrane in co-culture with astrocytes in the bottom chamber; the integrity of the bEnd.3 cell monolayer was assessed by using trans endothelial electrical resistance in an Epithelial Volt/Ohm meter equipped with STX2 electrodes (EMD Millipore Corporation). Uptake of MEs labeled with FM 4-64 and RNA cargos labeled with ExoGlow-RNA^TM^ was quantified by using a microplate fluorescence reader (BioTek Instruments, Inc.) and confocal microscopy imaging respectively. The transfer of IRDye-labeled miR-34a across a bEnd.3 cell monolayer was measured in dual chamber assays by using an Odyssey^R^ imaging system (LI-COR, Inc.).

### ME distribution in regions of the mouse brain

We developed an exosomes and cargo tracking (ECT) mouse on the C57BL/6J genetic background that enables studies of exosome and cargo trafficking among tissues, as well as studies of the transfer of ME from dam to pup [13]. Briefly, ECT mice express an open reading frame (ORF) coding for the exosome marker, CD63 [31] fused with enhanced green fluorescent protein (eGFP) flanked by loxP sites. In the presence of cre recombinase, the CD63-eGFP ORF is removed, and mice express an open reading frame coding for a fusion protein of CD63, near-infrared protein (iRFP), transmembrane domain and a second iRFP. The second iRFP localizes to the outer exosome surface and can be used to collect exosomes for cargo analysis.

Wild-type (WT) newborn C57BL/6J pups were fostered to ECT dams or WT dams from synchronized pregnancies and nursed for 17 days. Pups were euthanized and brains were fixed via transcardial perfusion of 4% paraformaldehyde, stored in phosphate-buffered saline and shipped to the Whole Brain Microscopy Facility at the University of Texas Southwestern Medical Center for analysis by serial two-photon tomography (STPT) [32; 33; 34; 35] and immunostaining of brain slices with anti-GFP antibodies (Invitrogen # A11122; 1:500 dilution). Eleven total brains from WT pups fostered to ECT dams across 3 separate litters and two brains from WT pups fostered to WT dams were used for STPT and anti-GFP immunostaining of isolated coronal brain sections. STPT is a high-resolution, high-throughput volumetric imaging strategy for assessing the regional distribution of native fluorescent labels throughout entire uncleared mouse brains via serial vibratome sectioning and mosaic two-photon imaging [32; 33; 34; 35]. Immunostained coronal brain sections were imaged using a Zeiss LSM 780 confocal microscope (Live Cell Imaging Facility, UTSW) at 20X and with the same acquisition parameters across samples. All animal studies in this paper were approved by the Institutional Animal Care Program at the University of Nebraska-Lincoln (protocols 1229 and 1713).

### Gene expression analysis

We assessed BME-dependent gene networks in the left hippocampus in male and female C57BL/6J mice (Jackson Labs., stock 000664). Briefly, C57BL/6 mice were fed ERD or ERS diets (**Box 1** [11; 36]) starting at age three weeks for seven weeks when mice were mated. Pups born to these breeders were continued on parental diets until age seven weeks. Pups were euthanized via transcardial perfusion with phosphate-buffered saline and brains were excised. The left hippocampus was dissected and flash frozen in liquid nitrogen for storage at -80°C. Total RNA was extracted by using miRNeasy Kit (Qiagen, Inc.) according to the manufacturer’s instructions, and the concentration, quality and integrity of RNA was analyzed as previously described [37; 38]. cDNA libraries were prepared by using a proprietary kit and samples were sequenced by using a paired-end 150 base-pair protocol and the NovaSeq platform (Illumina, Inc.) in the Beijing Genomic Institute. RNA-seq data were analyzed as previously described [15].

#### Box 1.

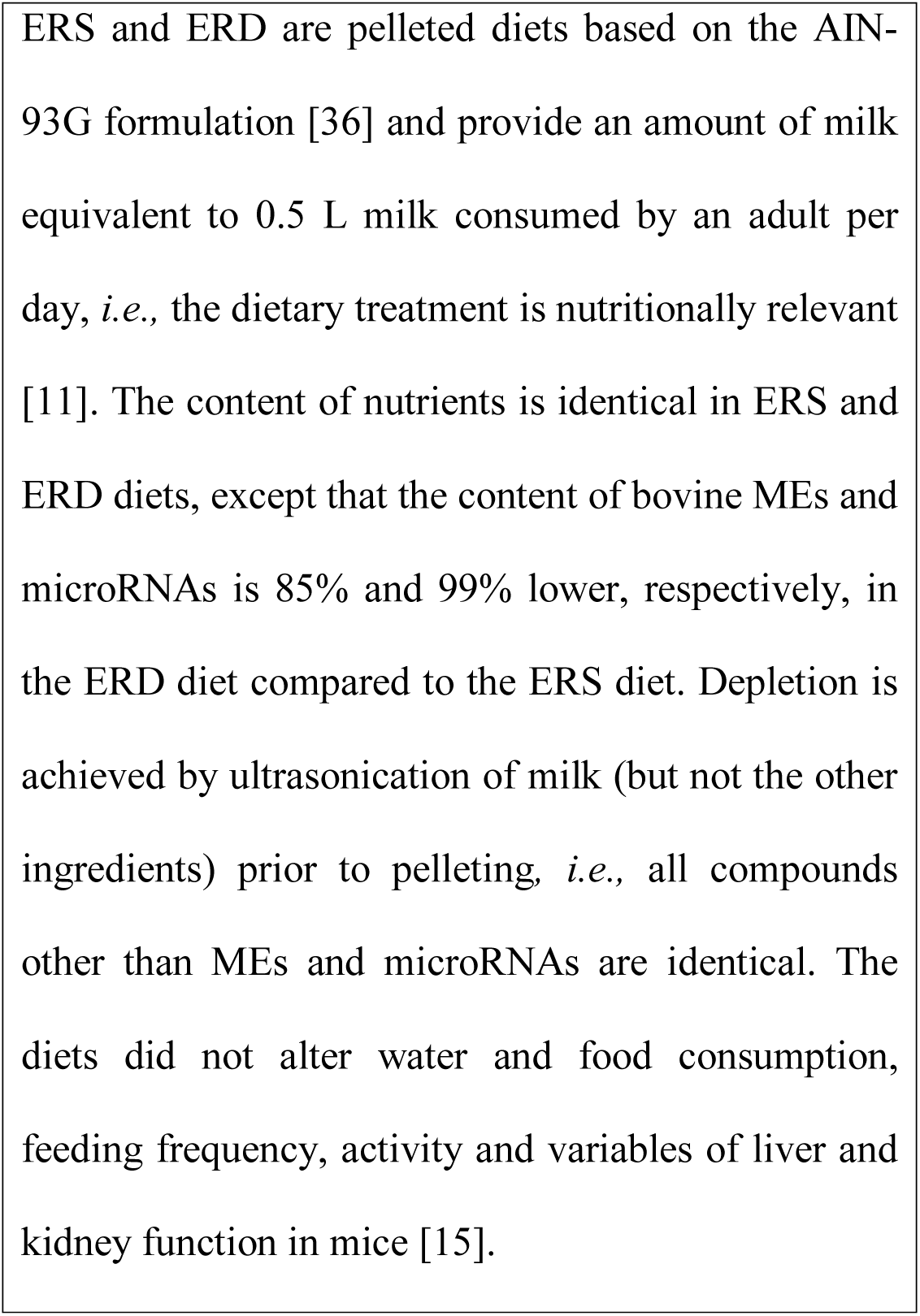
ERS and ERD rodent diets.

### Dendritic complexity of dentate granule cells

Neuronal dendrites are highly branched, tree-like structures, the morphological complexity of which is linked to signal integration and firing pattern of individual neurons and the functionality of neural circuitry [39]. Mice were fed ERD and ERS diets as described above. Hippocampal neurons from the left hemispheres of mice were stained using the Golgi-Cox method as previously described [40; 41]. Brightfield images of dentate granule cells in the suprapyramidal blade were collected in the Z plane at 20x magnification in 0.3-μm steps with an Olympus IX-81 inverted spinning disk confocal microscope using MetaMorph Advanced software version 7.1 (Molecular Devices). Three-dimensional dendritic structures of 3 – 5 granule cells were manually traced and reconstructed in each hippocampus using NeuroLucida version 2019.1.2 (MBF Bioscience). Dendritic complexity and other morphological features were quantified via Sholl analysis through NeuroLucida Explorer version 2019.1.2 (MBF Bioscience). Research staff was blinded regarding the treatment of mice.

### Phenotyping studies

Phenotyping studies focused on the assessment of spatial learning and memory (SLM), kainic acid-induced seizures, acoustic startle response and prepulse inhibition. These endpoints were chosen because the hippocampus has been implicated in SLM, kainic acid-induced seizures and acoustic startle response in mice and rats [42; 43; 44]. The choice of phenotypes is also consistent with our observations that MEs accumulated in the hippocampus (in addition to other regions) and altered the expression of genes implicated in axon guidance and calcium signaling in murine hippocampi in mice; dietary depletion of MEs led to a decrease in neuronal branching in murine dentate granule cells (see Results).

SLM was assessed at two ages by using the Barnes maze [45]. The Barnes maze measures the ability of a mouse to learn and remember the location of an escape hole on a circular surface with the help of a visual cue; low values represent strong test performance [45]. SLM experiments were conducted using the same mice that were subsequently used in RNA-sequencing analysis, except that additional mice were included in tests of SLM. SLM was also assessed in adult mice ages 12 – 15 weeks fed ERD and ERS diets starting at age 3 weeks. Mice were randomly assigned to diet groups and both sexes were studied.

In tests of seizure severity, C57BL/6J mice ages three weeks were fed ERD or ERS diets for 18 weeks when seizures were triggered by subcutaneous administration of kainic acid (25 mg/kg body weight) [46]. Kainic acid is a non-hydrolysable glutamate analog that binds to five glutamate receptors in the brain [47; 48], thereby causing neuronal excitotoxicity and seizures [46]. The severity of seizures was scored using a modified Racine scale [49; 50]. In the modified Racine scale, seizures are scored at timed intervals for two hours and the highest score in each 5-minute block are reported: 0, no seizure; 1, immobility; 2, forelimb and/or tail extension; 3, automatisms; 4, forelimb clonus, rearing, and/or falling; 5, repetition of stage 4; 6, tonic–clonic seizures; and 7, death.

Acoustic startle response and pre-pulse inhibition were assessed as previously described using the mice from the Barnes maze experiments one day after conducting the studies in the maze [51; 52]. Pre-pulse inhibition of the acoustic startle response was measured using prepulses of 68, 74 and 80 dB for 10 milliseconds with 65-dB background white noise, followed by a pulse of 105 dB for 20 milliseconds; intervals between stimuli were random 10 – 30 milliseconds. The percent prepulse inhibition of acoustic startle response was calculated as ((1 – (startle response at 105 dB with prepulse stimuli)/startle response for startle response at 105 dB without prepulse)) x 100. The acoustic startle response was scored ten times per mouse for each of the following conditions: 105-dB startle stimulus without prepulse, 105-dB startle stimulus with prepulses of 68, 74, 80 dB, and no startle stimulus with pulses of 80 dB. Test were performed by using an SR-LAB Startle Response System (San Diego Instruments, San Diego, California, USA).

### Statistical analysis

The F-test was used to assess the homogeneity of variances [53]. Some variances were heterogeneous, *e.g.,* Racine scale scores. Log transformation of these data resulted in homogenous data variation. Data from time-dependent transwell studies were analyzed by using repeated measures one-way ANOVA followed by Dunnett’s multiple comparisons test.The Kruskal-Wallis test was used for the analysis of data from acoustic startle response experiments. Data from both seizure studies and the Sholl analysis were analyzed by using repeated measures ANOVA and mixed procedure. The distance to soma was used as the within-subjects repeated measure. The model includes treatment as the fixed effect and mouse nested in treatment as the random term. Data analysis was conducted by using SPSS 27, SAS 9.4 and GraphPad Prism 9.0. *P* < 0.05 was considered statistically significant. Data are reported as mean ± SEM.

## RESULTS

### Transport of MEs by brain cells

bEnd.3 cells internalized BMEs by using a saturable process and secreted microRNA cargos across the basal membrane. The uptake of FM4-64-labeled BMEs was modeled using the Michaelis-Menten equation (**Fig. 1A**): transporter capacity (maximal velocity, V_max_) = 0.77 ± 0.18 x 10^11^ BMEs/(10,000 cells x 45 min) and affinity (Michaelis-Menten constant, K_m_) = 1.8 ± 2.0 x 10^11^ BMEs/mL). All subsequent studies in bEnd.3 cells were conducted under conditions when BME concentrations (6 x 10^11^ BMEs/mL) and incubation times (45 min) do not limit BME uptake (**Supplemental Fig. S2**). Z-stack confocal imaging confirmed that bEnd.3 cells internalized ExoGlow-RNA^TM^ labeled BMEs, as opposed to BMEs adsorbing to the cell surface (**Supplemental Fig. S3**). Upon internalization, the ExoGlow-RNA^TM^ labeled BMEs localized to the cell cytoplasm (**Supplemental Fig. S4**).

**FIGURE 1.**
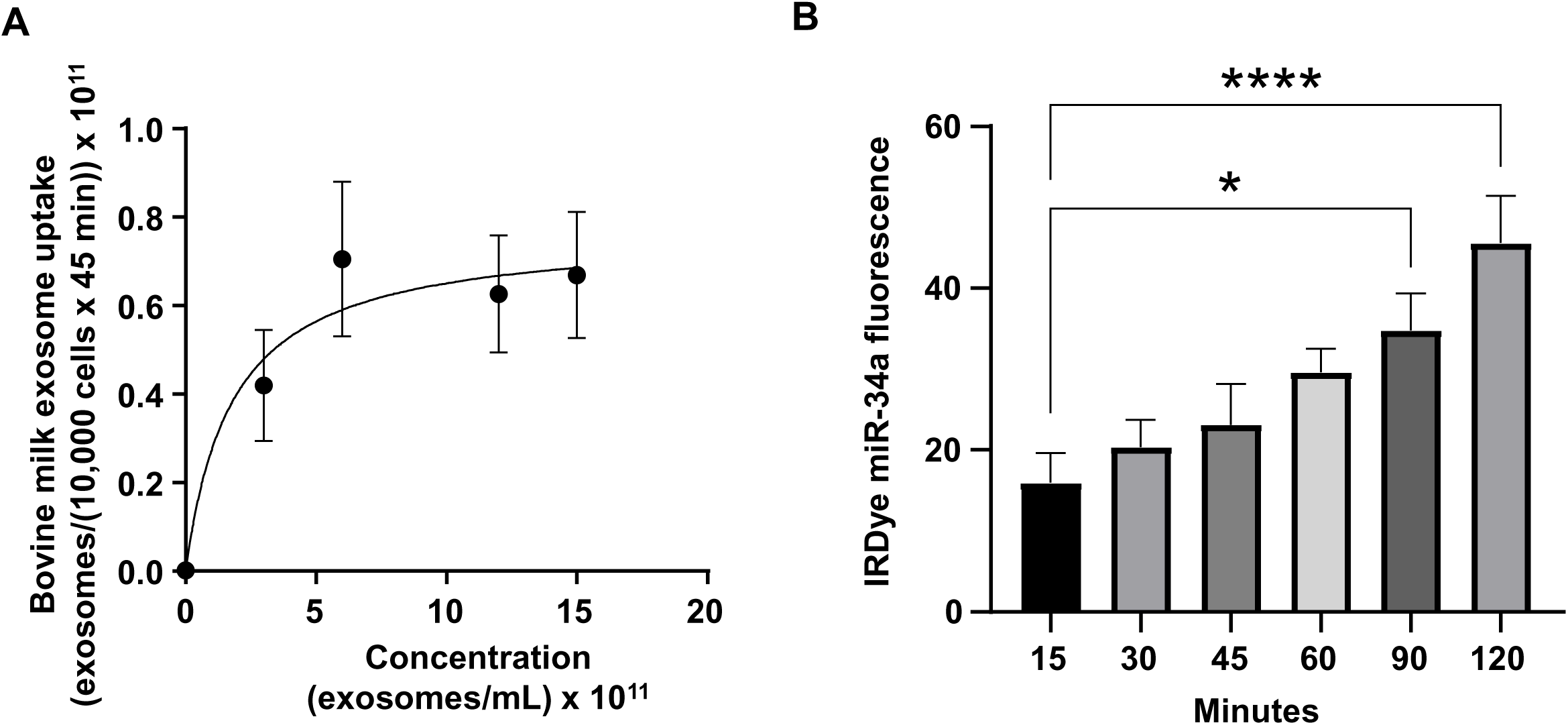
Transport of BMEs by murine brain cells. (A) Saturation kinetics of BME uptake by bEnd.3 cells. (B) Secretion of IRDye-labeled miR-34a, loaded into BMEs, in a dual chamber assay. Values are means ± SEMs, *n* = 3. \**P* < 0.05; \*\*\*\**P* < 0.001.

In a dual chamber model of transfer across the BBB, bEnd.3 cells secreted IRDye-labeled miR-34a, loaded into MEs, across the basal membrane into the bottom chamber in co-cultures with astrocytes (**Fig. 1B**). The integrity of the bEnd.3 cell monolayer on the semiporous membrane was assessed by measuring the trans endothelial electrical resistance and reached a plateau approximately four days after seeding the cells (**Supplemental Fig. S5**); absence of astrocytes in the bottom chamber caused a loss of monolayer integrity. BMEs that crossed the BBB, were internalized by brain cells, using brain macrophages, BV2 microglia as model. BME uptake by BV2 microglia followed saturation kinetics (**Fig. 2**): V_max_ = 0.66 ± 0.14 x 10^11^ BMEs/(10,000 cells x 45 min) and K_m_ = 1.9 ± 1.9 x 10^11^ BMEs/mL.

**FIGURE 2.**
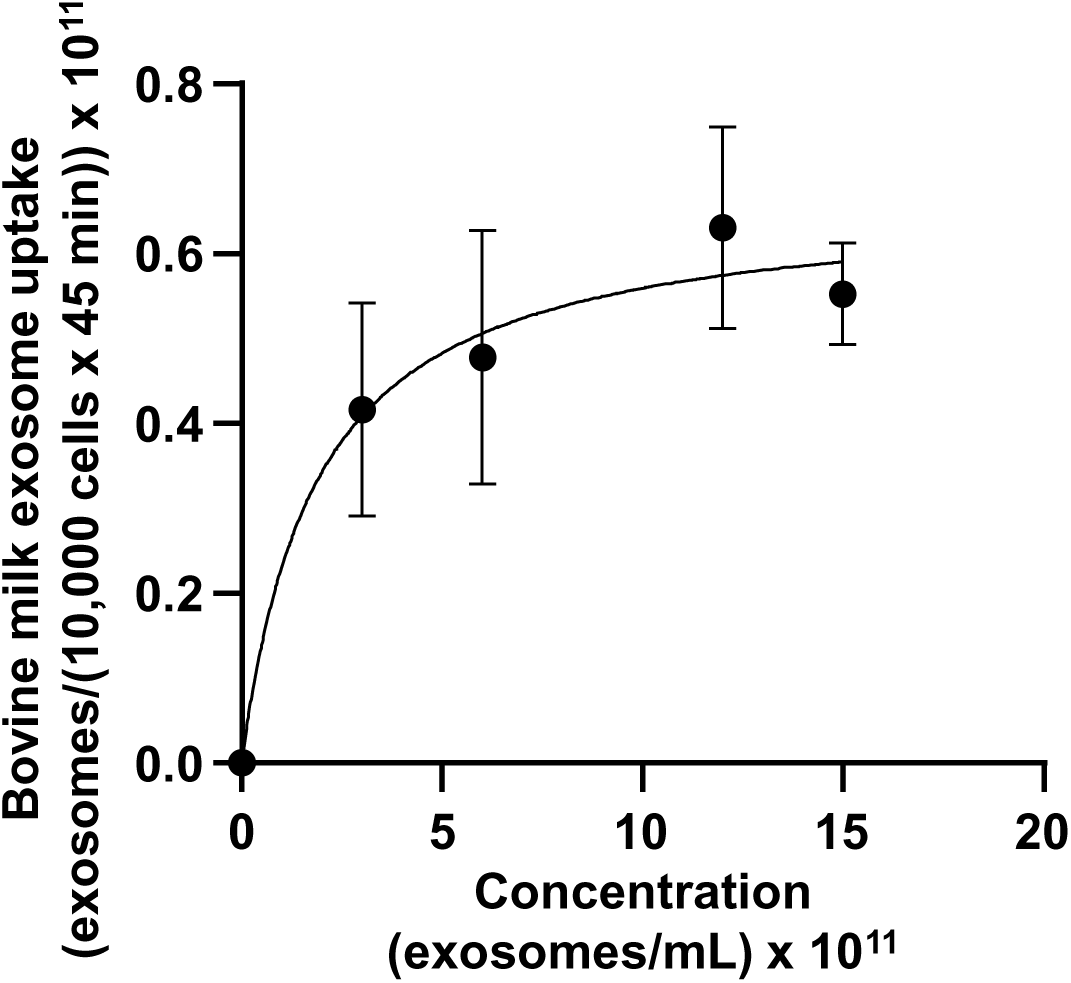
Transport of BMEs by murine BV2 microglia. Values are means ± SEMs, *n* = 3.

### ME distribution in the brain

eGFP-positive MEs accumulated primarily in the brain but also in the liver and small intestinal mucosa in WT pups fostered to ECT dams (**Fig. 3**). No eGFP fluorescence was detected in the the brain and liver in WT pups fosterd to WT dams, and the signal in the small intestine represents autofluorescence (control). Brains from WT pups fostered to either ECT or WT dams were subjected to serial two-photon tomography and anti-GFP immunostaining in isolated coronal brain sections. Accumulation of GFP labeled exosomes was evaluated in cortex, hippocampus and cerebellum using confocal imaging. Out of 11 brains from pups fostered to ECT dams across three litters, we observed positive anti-GFP staining in at least one of these three regions in seven of the brains. In total, brains from seven male pups and 4 female pups fostered to ECT dams were imaged. 2/7 male and 2/4 female brains did not show positive GFP signal. GFP signal was not observed in two brains from WT pups fostered to WT dams (one male and one female). Representative 2D coronal section images at the level of the dorsal hippocampus from the brain of a WT pup fostered to an ECT dam (**Fig. 4A**) and that of a WT pup fostered to a WT dam (**Fig. 4B**) shows robust accumulation of GFP labeled exosomes throughout the section from the brain of the pup which was fostered to the ECT dam (**Fig. 4A**), but no GFP signal was observed in the pup fostered to the WT dam (**Fig. 4B**). 3D renderings of the hippocampus (**Fig. 4C**) and the entire brain (**Fig. 4D**) from the pup fostered to the ECT dam demonstrate robust accumulation of GFP labeled exosomes throughout the hippocampus and many other brain regions. **Figures 4E-G** show confocal imaging of anti-GFP immunostaining in the brain of a different WT pup fostered to an ECT dam in the hippocampus, cerebellum and cortex. Positive GFP labeling was observed in each of these three brain regions. In contrast, no GFP labeling was seen in brains of pups fostered to WT dams as shown in **Figure 4H**.

**FIGURE 3.**
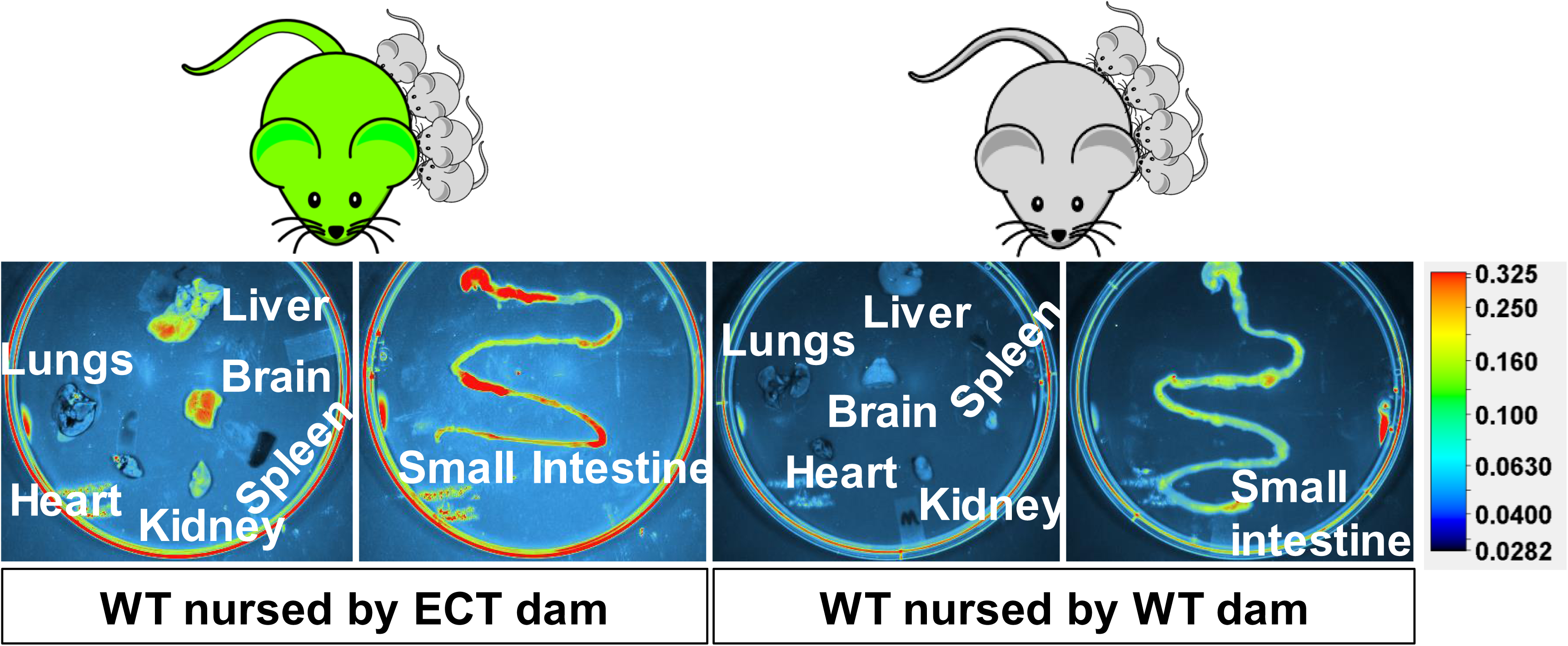
Accumulation of eGFP-positive milk exosomes in peripheral tissues and the small intestinal mucoca in wild-type pups fostered to ECT dams and nursed for 17 days. Wild-type pups fostered to wild-type dams served as controls.

**FIGURE 4.**
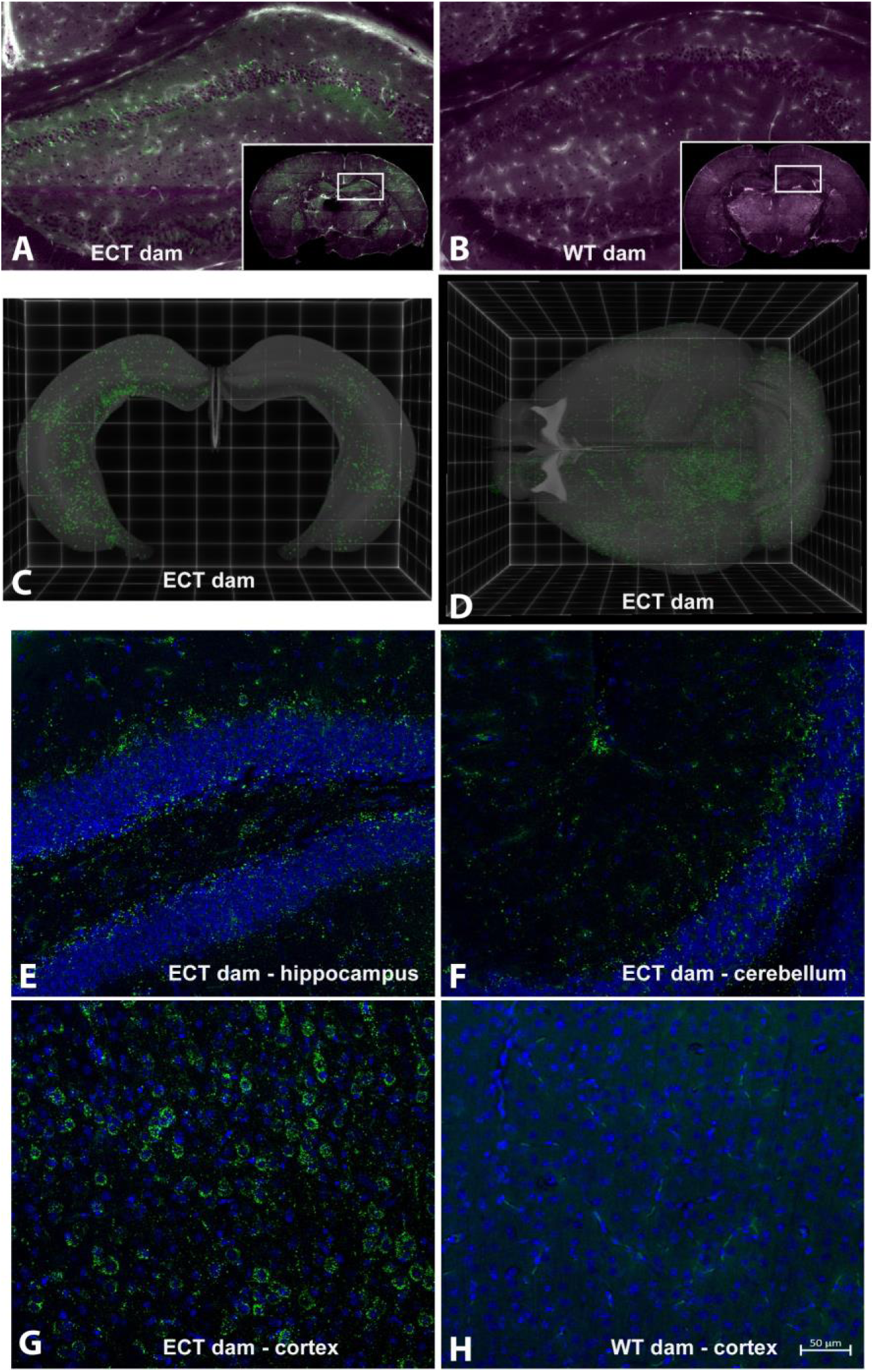
Serial Two-Photon Tomography (STPT) and confocal imaging in brains from wild-type pups fostered to either ECT or wild-type dams. (A) Single 2D section at the level of the dorsal hippocampus from a wild-type mouse pup fostered to an ECT dam acquired using STPT. Inset, whole coronal section with right hippocampus indicated with white box. Accumulation of GFP positive exosomes (green) is apparent in hippocampus and other areas throughout the section. Tissue autofluorescent signal is shown in magenta. (B) Single 2D section at the level of the dorsal hippocampus from a wild-type mouse pup fostered to a wild-type dam acquired using STPT. Inset, whole coronal section with right hippocampus indicated with white box. Accumulation of GFP positive exosomes was not detected. Tissue autofluorescent signal is shown in magenta. (C) 3D rendering of bilateral hippocampal volumes from a wild-type mouse pup fostered to an ECT dam acquired using STPT. Native GFP signal indicative of exosome accumulation is present throughout the entire hippocampal volume. (D) 3D rendering of entire brain volume from a wild-type mouse pup fostered to an ECT dam acquired using STPT. Native GFP signal indicative of exosome accumulation is present in many regions throughout the brain. (E-H) Confocal images from isolated coronal sections immunostained with anti-GFP antibodies, shown in green, and DAPI as nuclear counterstain, shown in blue. Images in Panels E, F, and G are from sections from the brain of a wild-type mouse pup fostered to an ECT dam and show accumulation of GFP positive exosomes in the hippocampus, cerebellum and cortex, respectively. Panel H shows an image from the cortex of the wild-type mouse pup fostered to a wild-type dam and no GFP signal was observed. Scale bar in H applies to Panels E-H.

### BME-dependent biological pathways

Dietary intake of BMEs altered gene expression in the brain. Two hundred ninety-five genes were differentially expressed in the left hippocampus in male mice fed ERD or ERS diets for seven weeks (**Fig. 5A**). Three genes (*Dcn*, decorin; *Nos1*, nitric oxide synthase 1; *Ndn*, necdin) was differentially expressed in females. Raw sequencing data can be accessed in the NCBI-BioProject database through accession number PRJNA783128. KEGG pathway analysis revealed 45 BME-dependent biological pathways in males, each with at least three BME-dependent genes (**Fig. 5B**). Pathways of calcium signaling and axon guidance were among the three top-ranked pathways. No KEGG pathways emerged in the analysis of mRNA expression data in females.

**FIGURE 5.**
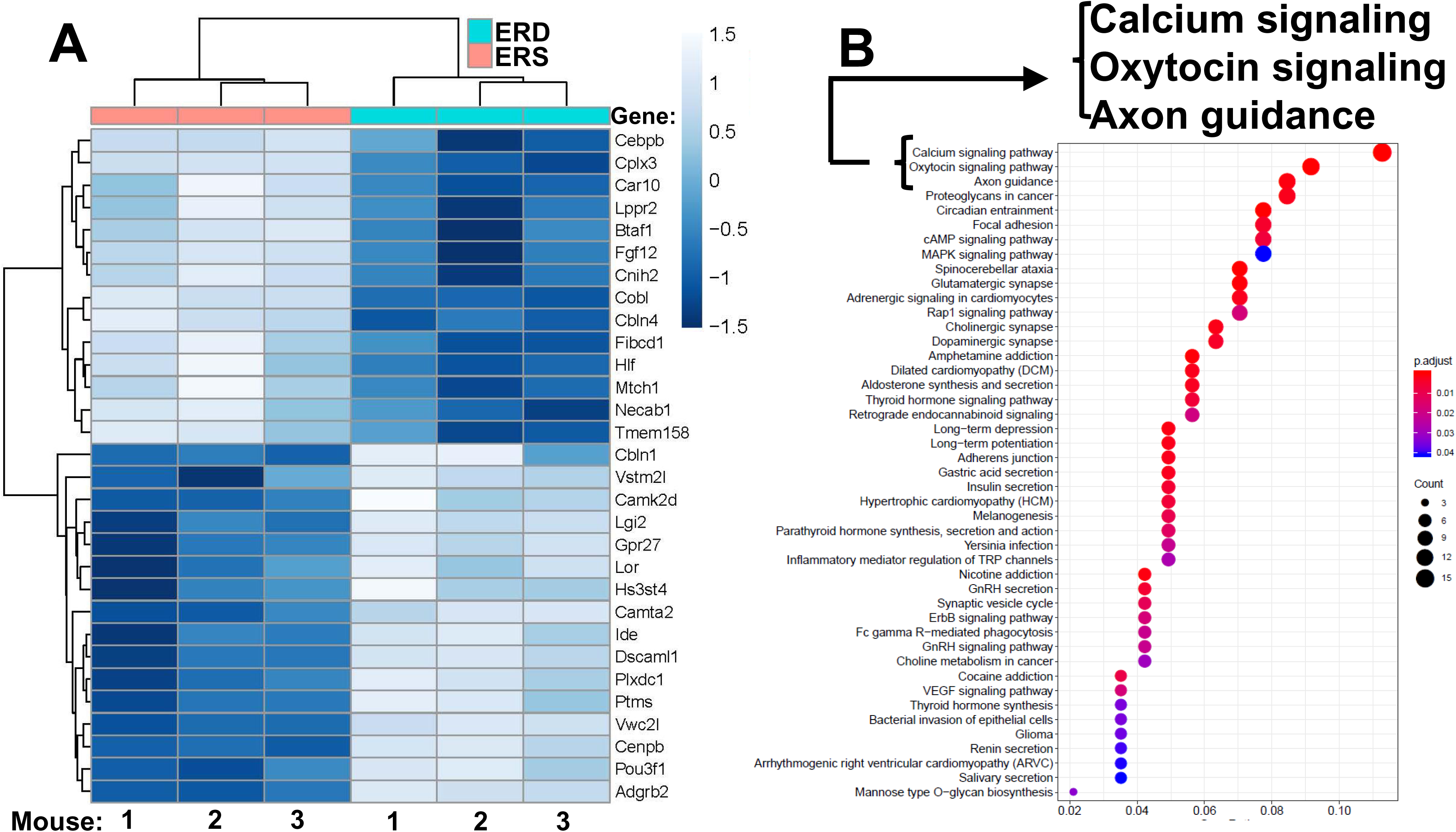
Gene expression. (A) Top 30 differentially expressed mRNAs in the left hippocampus in male pups. Means without a common letter differ (*P* < 0.05; *q* < 0.3; *n* = 3). (B) KEGG pathways. KEGG, Kyoto Encyclopedia of Genes and Genomes.

### Phenotypes of BME depletion

BME-dependent changes in gene expression and neuronal growth were associated with impaired SLM and increased the severity of kainic acid-induced seizures in mice fed the ERD diet compared to mice fed the ERS diet. For example, when female pups were nursed by dams fed the ERS diet for three weeks and continued on the maternal diet for one week, the mice performed 9 times better in the Barnes maze test of SLM compared to mice nursed by ERD dams and fed the ERD diet for one week (**Fig. 6A**). Diet effects on SLM were more modest in males than in females and in mice older than 4 weeks (**Supplemental Fig. S6**); the only exception are females ages 7 weeks in which diet effects were similar to females ages 4 weeks. Beneficial effects of BMEs on brain function were not limited to SLM but extended to seizure activity which was 5 times higher in male mice fed the ERD diet 20 minutes after administration of kainic acid compared to ERS controls (**Fig. 6B**, **Supplemental Table S1**). Note that the effect of diets on kainic acid-induced seizure activity were also detectable in females but effects were modest when compared to males and not statistically significant (**Supplemental Table S1**). Dietary effects on pre-pulse inhibition of acoustic startle response were modest and not statistically significant in most age groups and both sexes (data not shown).

**FIGURE 6.**
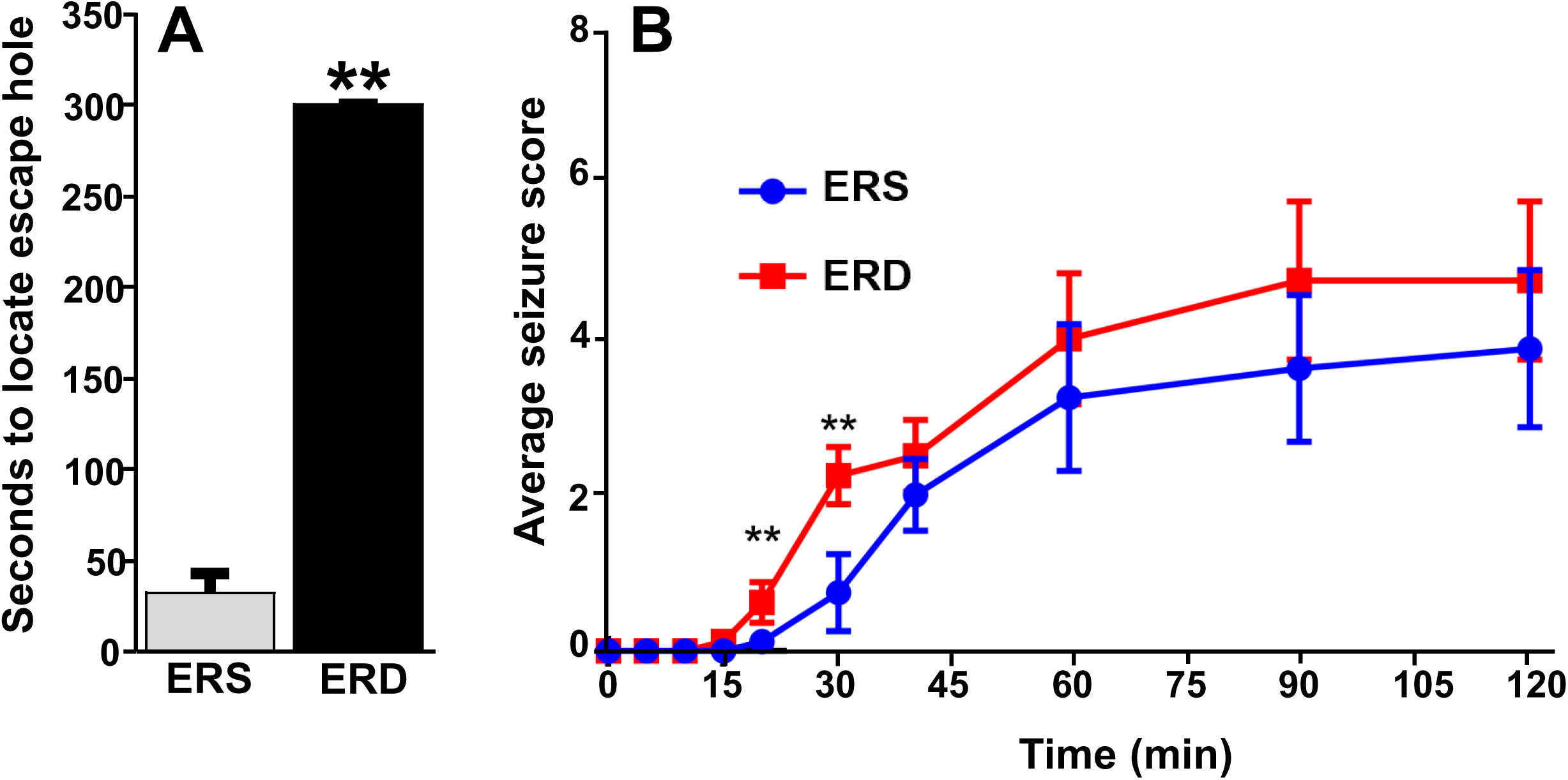
Effects of BME-defined diets on brain function in mice. (A) SLM in female pups, age 4 weeks. Values are means ± SEMs (\*\**P* < 0.01, *n* = 5). Means without a common letter differ. (B) Kainic acid-induced seizure activity in male mice ages 21 weeks. Values are means ± SEMs ( \*\**P* < 0.01, *n* = 8).

### Dendritic complexity

Given the preliminary evidence of neuronal accumulation of BME-derived microRNAs, as well as alterations in hippocampal pathways implicated in neuronal development and cognitive deficits in SLM after dietary BME depletion, we evaluated its impact on dendritic complexity of dentate granule cells in the hippocampus. Representative samples of traced three-dimensional dendritic architecture of dentate granule cells were shown in **Fig. 7A and 7B**. Sholl analyses evaluated dendritic complexity by quantifying the number of interactions between dendrites and soma-oriented concentric spheres of increasing diameters (**Fig. 7C**). There is a significant main effect of diet on dendritic complexity. This is evidenced by the smaller number of dendritic intersections in DG neurons from mice fed the ERD diet compared to mice fed the ERS diet (*P* < 0.05), suggesting that deficiency in milk exosome resulted in underdevelopment of neuron dendritic architecture in the developing brain. This difference was primarily due to a greater number of branch nodes of granule cells in ERS mice compared to ERD mice (**Table 1**). In addition, there was a trend towards a greater number of dendritic tips in ERS mice compared to ERD mice (*P* = 0.08). The number of primary dendrites and total dendritic length of dentate granule cells were not significantly different between diet groups (**Table 1**).

**FIGURE 7.**
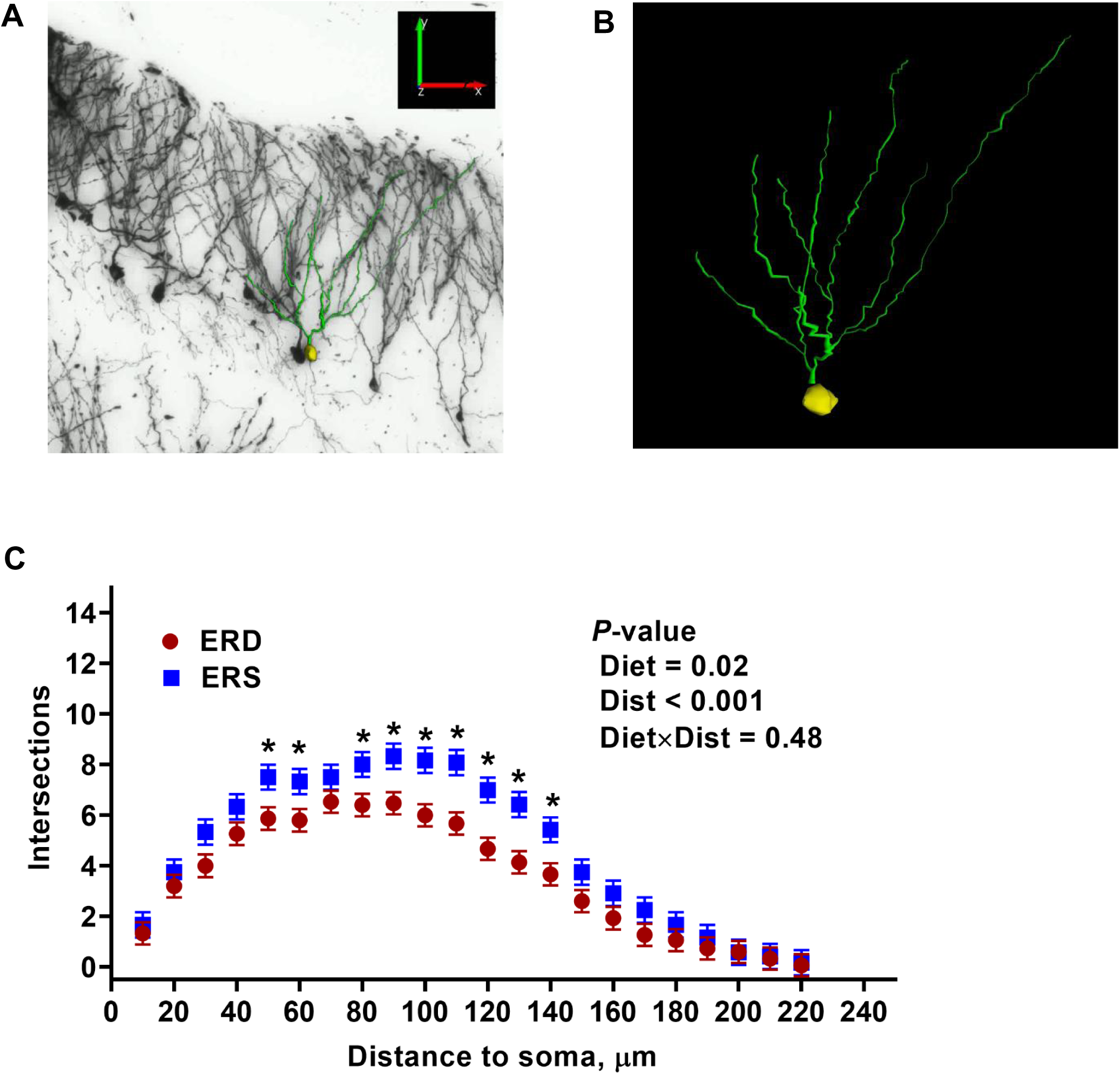
Three-dimensional dendritic architecture (A, B) and Sholl analysis of dendritic complexity (C) of dentate granule cells from murine hippocampus (*n* = 4 – 5 mice; 3 – 5 granule cells per mouse).

**Table 1.**
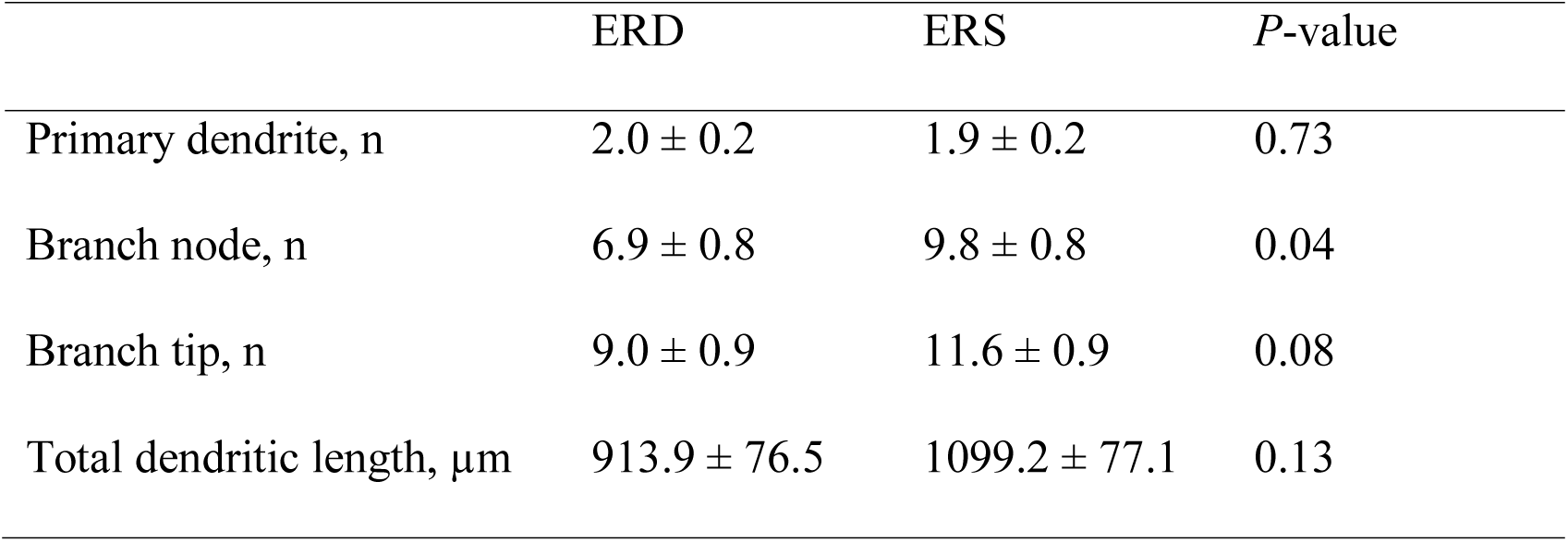
Effects of MEs on dendritic morphology of dentate granule cells in murine hippocampus (mean ± SEM).

## DISCUSSION

This is the first report that orally ingested MEs are transported across the BBB and accumulate in distinct regions of the brain. This report also provides experimental evidence that MEs deliver messages that alter gene expression and promote neuronal growth in the brain, and dietary depletion of MEs and cargos elicits phenotypes such as impaired SLM and increased severity of kainic acid-induced seizures. These discoveries are of great significance in infant nutrition because of the substantially greater content of MEs and microRNA in human milk compared to infant formulas, and milk is the sole source of nutrition in the first stages of mammalian life [23; 24]. Neurological phenotypes of ME depletion were evident in mice in this study, but causal relationships between the consumption of ME-poor formulas and ME-rich human milk in the neurological development of infants have yet to be investigated. There is circumstantial evidence in support of the theory that MEs might contribute towards optimal brain development in infants, although the studies were not designed to look specifically at MES and, therefore, other compounds in human milk could also have contributed to the positive effects of breastfeeding. For example, breastfed infants scored higher than formula-fed infants in tests of mental and psychomotor development, and breastfeeding increased white matter, sub-cortical gray matter volume and cortical thickness in infants [25; 56]. Breastfed infants scored higher in cognition tests than formula-fed infants in a meta-analysis [57]. To put the brain phenotypes reported in this study in context, when mice were fed n-3 polyunsaturated fatty acid (PUFA)-defined diets known to improve cognitive function, mice fed an n-3 PUFA-sufficient diet performed 1.35-fold better in the Barnes maze compared to mice fed an n-3 PUFA-deficient diet [58], whereas in our study, mice fed the ERS diet performed nine times better than the mice fed the ERD diet on the Barnes maze.

Confidence in the data reported here is high, because MEs accumulated and altered gene expression and dendritic architecture in the hippocampus, which is implicated in SLM, kainic acid-induced seizures and PPI of the ASR in mice [42; 44]. Also, there is a degree of specificity to the neurological phenotypes associated with ME depletion. For example, effects of BME depletion on muscle grip strength were modest in previous studies in mice and rats and this study revealed modest effects in rotarod and startle response tests [15; 19]. The exact mechanism of action by which BME depletion impairs SLM and increases the severity of kainic acid-induced seizures remains elusive. If neurological phenotypes are caused by a depletion of microRNA cargos, then miR-30d and let-7b might be the prime candidates for facilitating the phenotypes. MiR-30d and let-7b are the two most abundant microRNAs in human MEs and loss of miR-30d and let-7b signaling impaired axonal outgrowth in early neuronal development [24; 59].

This study suggests that some neurological effects of BMEs depend on sex and age. For example, the number of BME-dependent genes in the hippocampus was 40 times greater in males than in females ages seven weeks, and the effect of BME depletion on SLM was 1.7 times stronger in males than females ages 15-18 weeks. As for age effects, examples include that BME depletion had a stronger effect on SLM in female than male mice ages four weeks whereas effects of BME depletion on SLM were stronger in male than female mice ages 15-18 weeks. There is precedent for effects of age on brain development, e.g., the rate of gray matter accumulation peaked one or two years earlier in female than male adolescents [60].

This report, in conjunction with previous studies of ME and microRNA cargo bioavailability and phenotypes of depletion, suggests that MEs and microRNA cargos meet the definition of bioactive compounds by the National Cancer Institute which is “A type of chemical found in small amounts in plants and certain foods (such as fruits, vegetables, nuts, oils, and whole grains) which has actions in the body that may promote good health.” [61]. [62] Future lines of investigation will further delineate the roles of MEs and their miRNA cargos in neurodevelopment. For example, it will be important to determine whether our findings in mice translate into human populations, particularly infants. One could consider assessing neurological function in cohorts of infants fed ME-poor formulas and ME-rich human milk. Such studies will inform stakeholders whether the addition of MEs to infant formulas warrants consideration. Along these lines it will be important to assess whether phenotypes of ME depletion in infancy persist post weaning. Future studies will also need to fill knowledge gaps as to what cargos in MEs elicit neurological phenotypes and what signaling compounds these cargos affect.

## Acknowledgements

The authors’ responsibilities were as follows—FZ: performed the experiments, analyzed the data, performed the statistical analysis and the manuscript revision; PE: performed the experiments, analyzed the data, performed the statistical analysis and the drafting of the manuscript; EM: performed the experiments, analyzed the data, and performed the statistical analysis; SS: contributed to the experiments; SN: performed the experiments, analyzed the data, and performed statistical analysis; HD: contributed to the experimental design; analyzed the data and interpreted the data; WL: performed the experiments; JC: contributed to the experimental design; analyzed the data, interpreted the data and performed manuscript revision; PJ: contributed to the experimental design, analyzed the data, interpreted the data and performed manuscript revision, DMOR: contributed to the experimental design; analyzed the data, interpreted the data and performed manuscript revision; JZ: contributed to the experimental design, wrote the manuscript and took responsibility for the final content; and all authors read and approved the final manuscript.

## Abbreviations

BBB: blood-brain barrier
BME: bovine milk exosome
ECT: exosome and cargo tracking
eGFP: enhanced green fluorescent protein
ERD: exosome and RNA-depleted
ERS: exosome and RNA-sufficient
iRFP: near-infrared protein
ME: milk exosome
ORF: open reading frame
PBS: phosphate-buffered saline
PUFA: n-3 polyunsaturated fatty acid

