## Supplemental Materials for "Milk Exosomes Cross the Blood-Brain Barrier in Murine Cerebral Cortex Endothelial Cells and Promote Dendritic Complexity in the Hippocampus and Brain Function in C57BL/6J Mice"

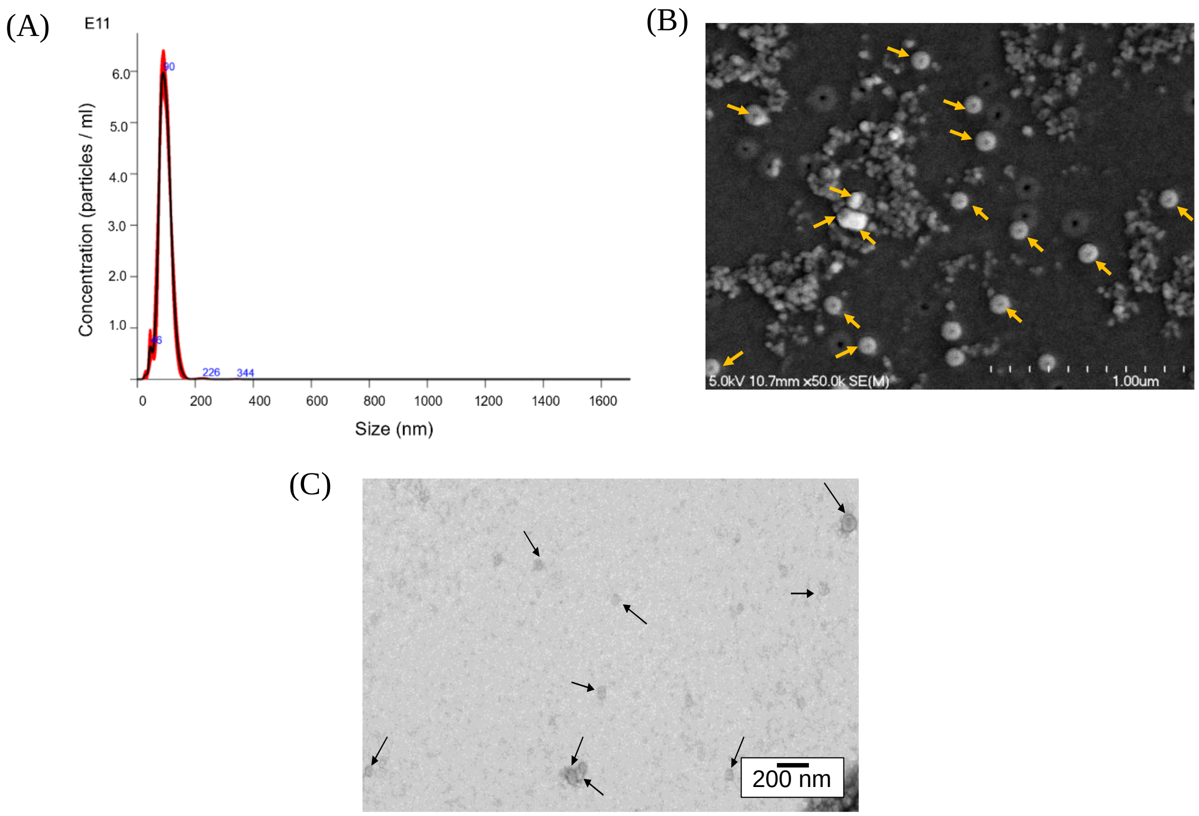


**Supplementary Fig. S1.** Bovine milk exosomes were authenticated by nanoparticle size analysis (A), scanning electron microscopy (B) and transmission electron microscopy (C).

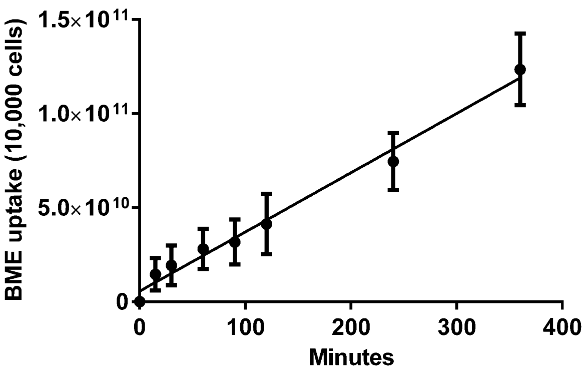


**Supplementary Fig. S2.** Time course of BME uptake by bEnd.3 cells. BMEs were labeled with FM4-64 ($n$ = 3).

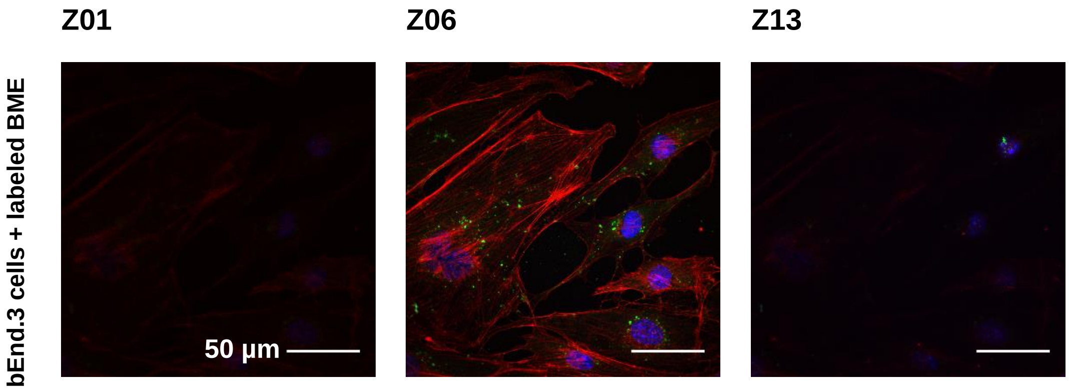


**Supplementary Fig. S3.** BME enter the interior of bEnd.3 cells. Images represent select focal planes acquired by Z-stack confocal microscopy. mRNA in BME was labeled with ExoGlow-RNA™ (green). Nuclei and actin (cytoplasm) were stained with DAPI (blue) and Alexa Fluor 568 phalloidin (red), respectively. Merged images are shown. Magnification = 60x.

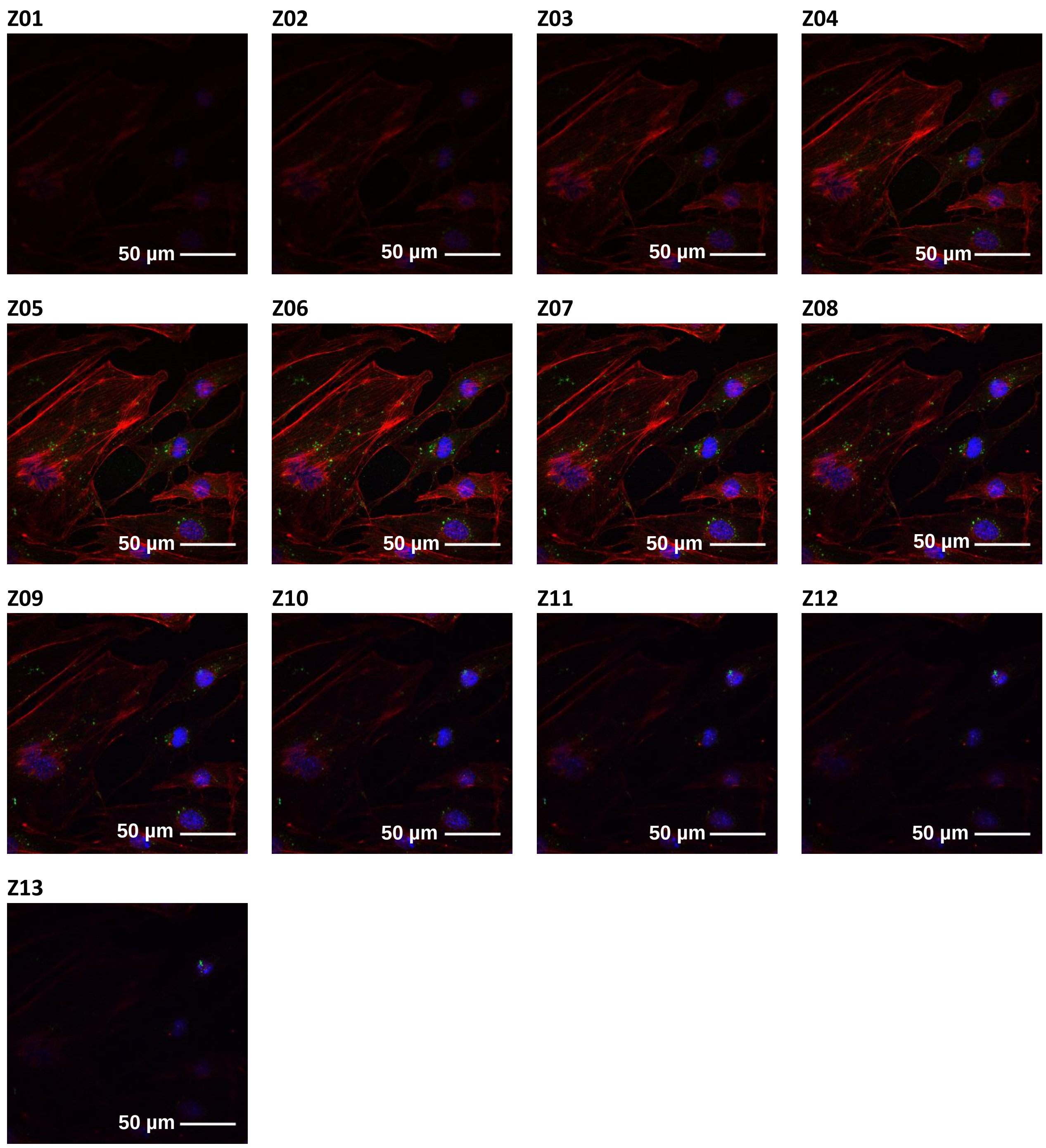


**Supplementary Fig. S4.** Overlay of individual Z-stack slices for bEnd.3 cells treatment group. Cells were incubated with BME in which mRNA was labeled with ExoGlow-RNA$^{TM}$ (green) for 24 h. Nuclei and actin (cytoplasm) were stained with DAPI (blue) and Alexa Fluor 568 phalloidin (red) respectively. Magnification = 60x.

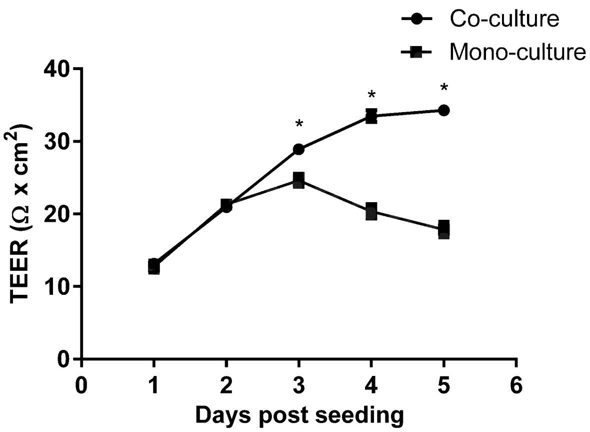


**Supplementary Fig. S5.** Daily transepithelial electrical resistance (TEER) measurements of bEnd.3 monocultures and co-cultures with C8-D1A astrocytes.

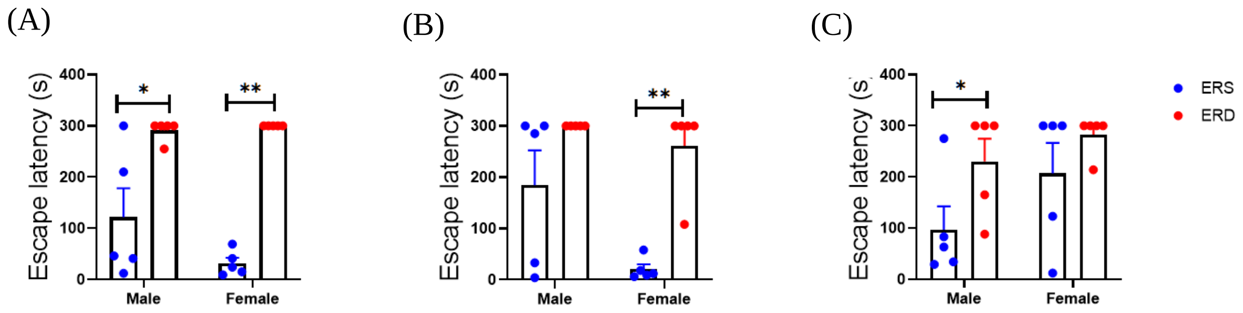


**Supplementary Fig. S6.** Effects of BME-defined diets on spatial learning and memory in C57BL/6J mice ages 4 weeks (A), 7 weeks (B) and 15-18 weeks (C). *$P$<0.05, **$P$<0.01 by Mann-Whitney test ($n$ = 5 per sex and age).

**Supplementary Table 1.** Racine scale scores of C57BL/6J mice ages 21 weeks fed exosome and RNA-defined diets starting at weaning.

| Diet (sex) | Score | | | | | | | | Mean[2] | SEM |
|---|---|---|---|---|---|---|---|---|---|---|
| | 0 | 1 | 2 | 3 | 4 | 5 | 6 | 7 | | |
| ERS (male) | 7 | 1 | | | | | | | 0.13 | 2.12 |
| ERD (male) | 4 | 3 | 1 | | | | | | 0.63** | 0.72 |
| ERS (female) | 1 | 3 | | | 1 | 1 | 1 | 1 | 3.13 | 0.30 |
| ERD (female) | | 2 | | 1 | 2 | 1 | | 2 | 4 | 0.22 |

[1]The revised Racine scale reports scores for the time after kainic acid administration when effects of treatment were largest (20 minutes for males, 120 minutes for females).

[2]Mean scores were calculated by multiplying the number of mice in each category with the numerical score in that category and dividing that number by the number of animals in the treatment group ($n = 8$). $^{**}P < 0.01$ by two-way ANOVA analysis (time *x* diet) of log transformed data.
